# An accelerated miRNA-based screen implicates Atf-3 in odorant receptor expression

**DOI:** 10.1101/001982

**Authors:** Shreelatha Bhat, Minjung Shin, Suhyoung Bahk, Young-Joon Kim, Walton D. Jones

**Affiliations:** Department of Biological Sciences, Korea Advanced Institute of Science and Technology (KAIST), Daejeon 305-701, Republic of Korea; School of Life Sciences, Gwangju Institute of Science and Technology (GIST), Gwangju 500-712, Republic of Korea

**Keywords:** *Drosophila*, genetic screening, miRNA, olfaction

## Abstract

Large scale genetic screening is tedious and time-consuming. To address this problem, we propose a novel two-tiered screening system comprising an initial “pooling” screen that identifies miRNAs whose tissue-specific over-expression causes a phenotype of interest followed by a more focused secondary screen that uses gene-specific RNAi. As miRNAs inhibit translation or direct the destruction of their target mRNAs, any phenotype observed with miRNA over-expression can be attributed to the loss-of-function of one or more target mRNAs. Since miRNA-target pairing is sequence-specific, a list of predicted targets for miRNAs identified in the initial screen serves as a list of candidates for the secondary RNAi-based screen. These predicted miRNA targets can be prioritized by expression pattern, and if multiple miRNAs produce the same phenotype, overlapping target predictions can be given higher priority in the follow-up screen.

Since miRNAs are short, miRNA misexpression will likely uncover artifactual miRNA-target relationships. Thus, we are using miRNAs as a tool to accelerate genetic screening rather than focus on the biology of miRNAs themselves. This two-tiered system allows us to rapidly identify individual target genes involved in a phenomenon of interest, often in less than 200 crosses. Here we demonstrate the effectiveness of this method by identifying miRNAs that alter *Drosophila* odorant receptor expression. With subsequent miRNA target prediction and follow-up RNAi screening we identify and validate a novel role for the transcription factor Atf3 in the expression of the socially relevant receptor Or47b.

## Introduction

The development of transgenic RNAi fly stock libraries (e.g., the Vienna Drosophila RNAi library (Dietzl et al., 2007) and the Transgenic RNAi Project (TRiP)) was a tremendous boon to the *Drosophila* community because they permit tissue-specific knockdown of almost all genes in the genome. These resources permit genome-wide screens for genes associated with almost any phenotype of interest. Unfortunately, the sheer size of these libraries—more than 22,000 stocks in the case of the Vienna library—means performing such screens remains labor-intensive and tedious. In this paper, we describe our development of a two-tiered screening protocol comprising an initial pooling screen using miRNA over-expression that generates a list of candidate genes involved in a phenotype of interest and a secondary screen using gene-specific RNAi that narrows this list of candidates to a single responsible target gene. We suggest that this protocol can accelerate the identification of novel genes involved in a broad range of phenotypes.

MicroRNAs are short, endogenous, single-stranded RNA molecules that act in the context of the miRISC protein complex to either inhibit translation or induce the degradation of target mRNAs (Bartel, 2004). Since the miRNA-target mRNA relationship is determined primarily by a short seed sequence at the 5’ end of each miRNA (Lewis et al., 2003; Lai, 2002), the complement of which may occur in multiple copies scattered over the genome, many miRNAs are capable of down-regulating multiple targets. The specificity of the relationship between a miRNA seed sequence and its complements in the open reading frames and 3’-untranslated regions (3’-UTRs) of target mRNAs spurred the development of bioinformatic algorithms that convert mature miRNA sequences into lists of potential mRNA targets (Rajewsky, 2006). These lists of candidate targets, however, are plagued by large numbers of false positives because the algorithms that generate them can fully account for neither the precise spatial and temporal patterns of miRNA and target mRNA expression nor target site availability. In other words, a miRNA may be capable of down-regulating a particular target and never actually do so, either because the two are never simultaneously expressed in the same tissue or because RNA-binding proteins or RNA folding render the target site inaccessible. It follows conversely that miRNA over-expression in arbitrary tissues using the binary GAL4/UAS expression system would likely lead to artifactual miRNA-target mRNA pairings. Rather than seeing such artifacts as a potential drawback of using a library of UAS-miRNA stocks, we propose that such pairings can be useful as part of a two-tiered screening system.

To this end, we generated a library of 131 UAS-miRNA fly stocks that permit tissue-specific over-expression of 144 *Drosophila* miRNAs. An ideal validation of our proposed screening process would not only identify novel genes, but also confirm the role of known genes. We identified an ideal proving ground for our two-tiered miRNA-based screening concept in the *Drosophila* olfactory system.

The olfactory sensory neurons (OSNs) of adult *Drosophila* are housed in porous hair-like sensilla that cover the paired antennae and maxillary palps. Olfactory sensilla are divided into 3 main classes by their shape and 17 subclasses by their odor response profile (Couto et al., 2005). The odor response profile of an OSN is determined by its expression of the obligatory olfactory co-receptor Orco and one or very few of the adult odor-specific odorant receptors (ORs) (Vosshall and Stocker, 2007). The spatial arrangement of the 17 subclasses of adult olfactory sensilla on the antenna, the arrangement of the OSNs themselves, the precise pattern of OR expression, and the wiring of the antennal OSNs into the appropriate glomeruli of the antennal lobe are all highly stereotyped from fly to fly, indicating tight developmental control of every step in the process.

Recently Jafari et al. reported the results of a large-scale RNAi screen that identified seven transcription factors, permutations of which determine the odorant receptor expressed by each population of olfactory neurons in the adult *Drosophila* antenna. Despite the success of their screen, Jafari et al. extrapolated from the complexity of the fly olfactory system and estimated that at least three more unidentified transcription factors are likely part of the combinatorial code that determines OR expression (Jafari et al., 2012).

In their screen, Jafari et al. combined the Pbl-GAL4 driver line, which is strongly expressed in peripheral sensory neurons including the antennal olfactory neurons, with a pair of OR promoter fusions (i.e., Or47b and Or92a) to a membrane-tethered GFP that act as reporters of OR expression. We obtained these lines and by crossing them to our library of UAS-miRNA stocks we were able to identify miRNAs whose over-expression eliminates Or47b expression, Or92a expression, or both. We chose to proceed with the miRNAs that affect Or47b expression (i.e., bantam, miR-2a-2, miR-33, miR-263a, miR-308, miR-973/974, and miR-2491). We then used existing bioinformatic tools to generate lists of their putative mRNA targets, compare the lists for overlap, and define a short list of candidate genes for a small follow-up RNAi screen. In this follow-up screen, we identified a previously unknown role for Activating transcription factor 3 (Atf3) in the expression of Or47b.

In addition, Eip93f, the only other transcription factor known to be involved in Or47b expression, is also a predicted target of some of the miRNAs that cause loss of Or47b expression. In other words, we were able to validate our screening system on a known transcription factor and rapidly (i.e., in a small number of genetic crosses) identify a previously unknown player in OR expression, Atf3.

## Results

### UAS-miRNA library generation

According to the most recent release of miRBase, roughly 65% of the 238 annotated miRNAs in the *Drosophila* genome are “high confidence” annotations (Kozomara and Griffiths-Jones, 2013). When we began this project, although there were no publicly available fly stocks for tissue-specific miRNA over-expression, Silver et al. had published a group of UAS-DsRed-miRNA DNA constructs representing many of these important miRNAs (Silver et al., 2007). In these constructs, 400-500 nucleotide pri-miRNA regions were inserted into the 3’-UTR of the fluorescent protein DsRed, itself part of a P-element-based UAS transgenesis vector. To facilitate the genetics of our proposed screening protocol, we decided to generate a library of site-specific insertions of these constructs. In addition to adding attB recombinase recognition sites to 44 of these UAS-DsRed-miRNA constructs, we amplified and inserted 400-500 nucleotide pri-miRNA sequences into the 3’-UTR of a similar UAS vector containing either DsRed or mCherry and an attB site. Clustered miRNAs were generally cloned en masse, but some were cloned individually (Fig. 1A). In total, our UAS-miRNA library includes 131 stocks covering 144 *Drosophila* miRNAs (see Supplemental Table S1 for details).

**Figure 1.**
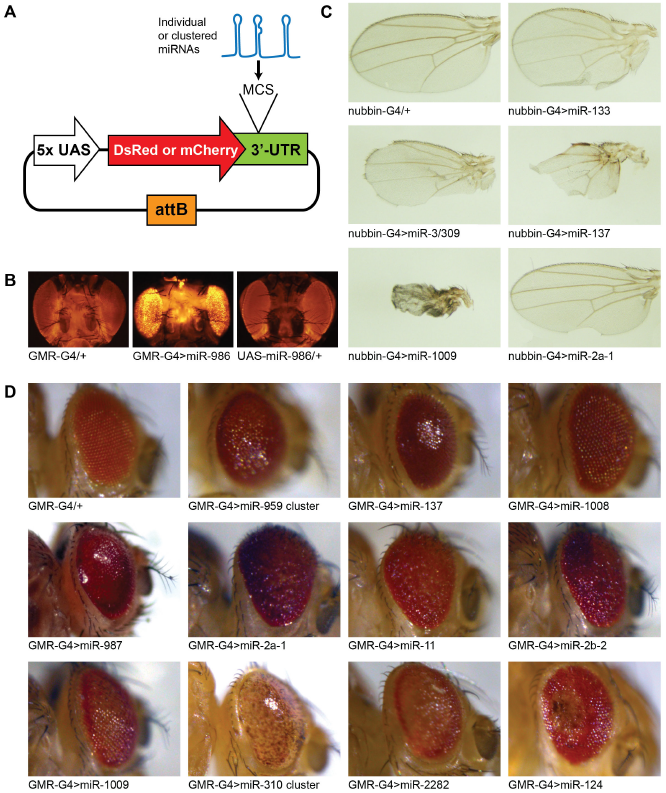
Generation and validation of a library of UAS-miRNA *Drosophila* stocks. **A.** Vector schematic for UAS-miRNA generation. All miRNAs (clustered and individual) were inserted into a multiple cloning site in the 3’-UTR of a fluorescent marker. **B**. DsRed fluorescence induced in the eye with GMR-GAL4 is a proxy for miR-986 expression (center). Heterozygous GMR-GAL4 and UAS-miR-986 controls (left and right, respectively) show only low levels of auto-fluorescence. **C**. miRNA expression in developing wings using nubbin-GAL4 induces hyperproliferation (miR-133), loss of marginal bristles and cross-veins (miR-3/309), severe malformations (miR-137 and miR-1009) as well as simple notching (miR-2a-1). **D**. GMR-GAL4 driven expression of several miRNAs induces rough, small, smooth, necrotic, or de-pigmented eyes.

### Verification of miRNA expression

We used four techniques to verify the expressibility and function of our UAS-miRNA lines. Since the miRNA sequences were placed in the 3’-UTR of either DsRed or mCherry, we were able to visually verify the expression of the marker after crossing it to the eye driver GMR-GAL4 (Fig. 1B). We also observed obvious developmental phenotypes induced by miRNA over-expression in both the wing and eye. Using the wing driver nubbin-GAL4, we observed miRNAs that induced wing notching, thinning of wing veins, loss of marginal bristles, and other more severe deformities (Fig. 1C). With GMR-GAL4, we observed rough eyes, smooth eyes, small eyes, and changes in pigmentation (Fig. 1D). We also directly verified the over-expression of a few select miRNAs using commercially available Taqman qPCR kits (data not shown). Since we began generating these UAS-miRNA lines, three other collections of UAS-miRNA lines using a similar cloning strategy were published and made available to the *Drosophila* community (Szuplewski et al., 2012; Bejarano et al., 2012; Schertel et al., 2012) highlighting their general usefulness.

### Two-tiered miRNA-based screening

The three collections of UAS-miRNA stocks published recently were presented as tools either for identifying novel miRNA functions (Bejarano et al., 2012; Schertel et al., 2012) or for use in the context of screens for modifiers of existing developmental phenotypes (Szuplewski et al., 2012). Rather than using our UAS-miRNA lines to study miRNA biology, we decided to examine whether they could also be used as a tool permitting a “pooling” pre-screen designed to limit the focus and thus accelerate a secondary, but more traditional RNAi-based loss-of-function screen. In such a scheme, after identifying miRNAs whose over-expression induces a phenotype of interest, bioinformatic target prediction provides a list of candidate genes for a follow-up RNAi screen. Since miRNAs inhibit the translation of or induce degradation of their target mRNAs, any phenotype observed with miRNA over-expression should be replicable via RNAi knockdown of the responsible target. Figure 2A presents a generalized flowchart of this two-tiered screening strategy.

**Figure 2.**
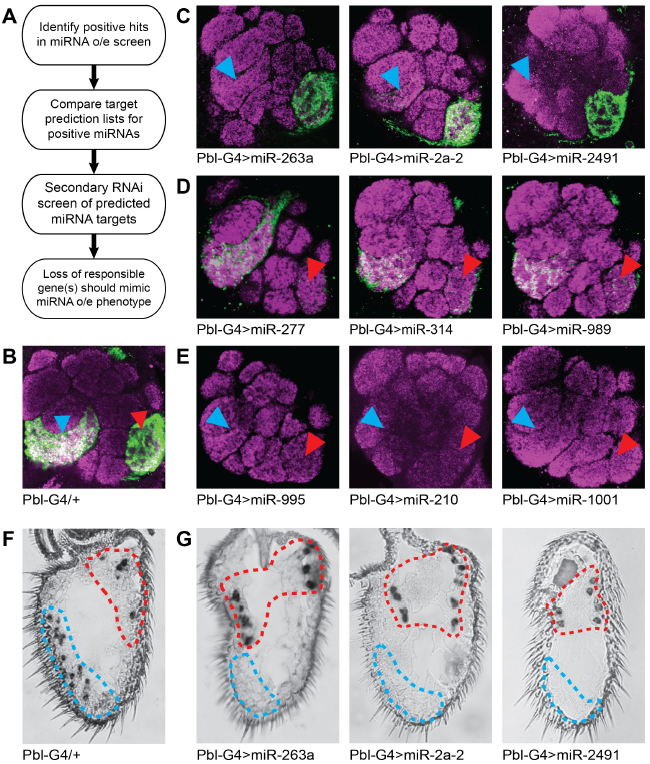
Primary screen for miRNAs whose over-expression alters OR reporter expression in the antennal lobe. **A.** A two-tiered, miRNA-based screening workflow can accelerate loss-of-function genetic screens by limiting the number of necessary gene-specific RNAi crosses. **B**. Antennal lobe (AL) staining of the control Pbl-GAL4/+; Or47b>CD8::GFP, Or92a>CD8::GFP/+ controls shows strong GFP staining in both the Or47b glomerulus VA1lm (blue arrowhead) and the Or92a glomerulus VA2 (red arrowhead). **C**. AL staining for 3 of the 7 miRNAs whose over-expression induces loss of the Or47b reporter (blue arrowheads). **D**. AL staining for 3 of the 4 miRNAs whose over-expression induces loss of the Or92a reporter (red arrowheads). **E**. AL staining for the 3 miRNAs whose over-expression induces loss of both the Or47b and Or92a reporters (blue and red arrowheads). **F**. Mixed probe *in situ* on the same control genotype as in B. Or92a-expressing cells are circled in red, while Or47b-expressing cells are circled in blue. **G**. Mixed Or47b/Or92a probe *in situs* on antennae of the genotypes shows in D confirm the loss of Or47b expression induced by miR-263a, miR-2a-2, and miR-2491.

### Primary screen for miRNAs that modify OR expression

As stated in the introduction, we settled on *Drosophila* olfactory neurons to demonstrate our miRNA-based screening strategy. Like Jafari et al., we chose Pbl-GAL4 to drive expression of UAS-miRNAs in adult olfactory neurons because it is expressed strongly in peripheral sensory neurons beginning 12-18 hours after pupation (Sweeney et al., 2007). By combining Pbl-GAL4 with fusions of two odorant receptor promoters (i.e., Or92a and Or47b) to the coding sequence for a membrane-tethered GFP and crossing in the UAS-miRNAs, we were able to visualize the effects of miRNAs on OR expression. Or92a is expressed in large basiconic sensilla in a class of OSN called ab1B, which respond to 2,3-butanedione (de Bruyne et al., 2001). Or47b is expressed in trichoid sensilla in a class of OSNs called at4, which respond to socially relevant fly-derived odors (van der Goes van Naters and Carlson, 2007). We chose Or92a and Or47b because they are expressed in olfactory neurons that innervate prominent, well-separated glomeruli in the antennal lobe (i.e., VA2 and VA1lm) and whose cell bodies are easily distinguishable upon antennal RNA *in situ* hybridization.

For the primary miRNA over-expression screen, we crossed a stable homozygous stock containing Pbl-GAL4 and the Or47b and Or92a promoter fusions to all of our UAS-miRNA lines. We observed that Pbl-GAL4 mediated miRNA over-expression for 20 of 131 UAS-miRNA lines is lethal or nearly lethal (see Supplemental Table 1). For the remaining miRNAs, we compared the levels of the GFP reporters of OR expression in the antennal lobes of flies over-expressing miRNAs with a heterozygous control (Fig. 2B). In doing so, we were able to divide miRNAs into three categories: those that reduce the intensity of the Or47b reporter in VA1lm (Fig. 2C, blue arrowheads), the Or92a reporter in VA2 (Fig. 2D, red arrowheads), or both (Fig. 2E).

We decided to focus our follow-up on the miRNAs that reduce the expression of the Or47b reporter because only one transcription factor, Eip93f, is associated with Or47b expression (Jafari et al., 2012). In addition, some of the miRNAs that cause loss of Or92a also cause a less significant loss of Or47b. In other words, the loss of Or47b phenotypes are less ambiguous. Before proceeding to miRNA target identification, we verified the miRNA-induced loss of Or47b using RNA *in situ* hybridization with mixed riboprobes against both Or47b and Or92a. All together we identified seven miRNA lines (Pictured: miR-263a, miR-2a-2, and miR-2491; Not shown: bantam, miR-33, miR-308, and miR-973/974) that reduce the expression of the Or47b reporter. We were able to verify that these lines eliminate Or47b (circled in blue) expression when combined with the Pbl-GAL4 driver while leaving Or92a (circled in red) expression intact (Fig. 2F,G).

### Secondary RNAi screen for miRNA targets that modify OR expression

According to the screening protocol outlined in Fig. 2A, we next compiled lists of possible target genes for each of the miRNAs whose over-expression eliminates Or47b expression. We initially compared the outputs of multiple target prediction algorithms (i.e., TargetScan, miRanda, and PITA), as this seems to be standard practice (Hyun et al., 2009; Silver et al., 2007). Figure 3A compares the numbers of candidate targets predicted for each miRNA that induces loss of Or47b by the TargetScan (Ruby et al., 2007), miRanda (Enright et al., 2003), and PITA (Kertesz et al., 2007) algorithms.

**Figure 3.**
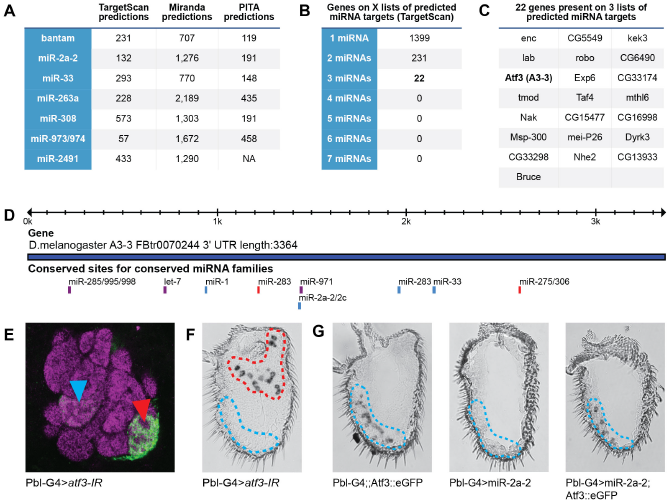
Secondary screen for regulators of Or47b expression using gene-specific RNAi. **A**. Tabular summary of the target predictions made by three different algorithms for each miRNA whose over-expression reduces Or47b expression. **B**. The number of genes that appear on the TargetScan candidate target lists in A. **C**. Twenty-two genes, including Atf3, appear on the candidate target lists for 3 different miRNAs. **D**. A schematized view of the Atf3 (A3-3) 3’-UTR showing the locations of seed matches for miR-2a and miR-33, which reduce Or47b expression. **E**. Atf3 knockdown in all olfactory neurons using Pbl-GAL4 reduces expression of an Or47b reporter, but spares expression of an Or92a reporter. **F**. Antennal *in situ* hybridization with mixed riboprobes that recognize Or47b (blue) and Or92a (red) reveals that Atf3 knockdown eliminates Or47b expression, but spares Or92a expression. **G**. An Atf3::eGFP fusion protein lacking the Atf3 3’-UTR, which contains the miR-2a-2 binding site, rescues the loss of Or47b expression observed with miR-2a-2 over-expression.

Typically, miRNA-based screens using GAL4 lines that drive expression in well-characterized tissues would allow a list of predicted miRNA targets to be prioritized based on expression in the tissue of interest. In this case, however, without access to transcriptome data for olfactory neurons, we decided to prioritize our secondary RNAi screen by looking for overlap between the lists of candidate targets for each miRNA whose over-expression eliminated Or47b expression. We decided to focus on TargetScan because it predicts miRNA seed matches in both 3’-UTRs and open reading frames as well as provides a way to query more recently identified miRNAs like miR-2491. PITA does not provide easy access to this option, and the miRanda algorithm produces too many predicted targets. TargetScan predicts a total of 1399 unique targets for the seven UAS-miRNA lines that eliminate Or47b expression, but only 231 unique candidates appear on two lists and 22 unique candidates appear on three lists (Fig. 3B,C).

We next screened through the list of candidate targets that appear on the prediction lists for multiple miRNAs by combining gene-specific UAS-candidate-IR (inverted repeat) lines with the control Pbl-GAL4; Or47b*>*CD8::GFP, Or92a*>*CD8::GFP genotype. In this secondary screen, knockdown of responsible targets can be expected to mimic the miRNA-induced loss of Or47b. This is how we were able to confirm a role for Activating Transcription Factor 3 (Atf3 or A3-3) in the expression of Or47b. Atf3 appears on the predicted target lists for three miRNAs whose over-expression eliminates Or47b expression (i.e., miR-2a-2, miR-33, and miR-2491). Figure 3D indicates the positions of the miR-2a-2 and miR-33 binding sites in the Atf3 3’-UTR. The miR-2491 binding site in the 3’-UTR of Atf3 is conserved across several species of *Drosophila*, but because it is not part of the standard TargetScan search parameters it does not appear in the exportable graphic used to make Figure 3D. It should also be noted that over-expression of several miRNAs (e.g., let-7, miR-285, etc.) with predicted binding sites in the Atf3 3’-UTR fails to eliminate Or47b expression, presumably due to issues with binding site accessibility. The fact that it is currently impossible to predict which of the clear miRNA seed match sites represent true positives is what makes a follow-up gene-specific RNAi screen necessary. Figure 3E shows the result of our secondary RNAi screen in which knockdown of Atf3 dramatically and specifically reduces the expression of the Or47b reporter (blue arrowhead) and not the Or92a reporter (red arrowhead). Furthermore, loss of Atf3 eliminates Or47b expression as assessed via mixed probe *in situ* hybridization while leaving Or92a expression unaffected (compare Fig. 3F to Fig. 2C). Since we are using miRNAs as a screening tool, the precise mechanisms by which they produce their phenotypes are not relevant to the results of the follow-up screen, but in the interest of completeness we confirmed that the loss of Or47b expression induced by miR-2a-2 over-expression is rescued by introduction of an Atf3::eGFP protein fusion that lacks its 3’-UTR and thus miR-2a-2 binding site (Fig. 3G).

As additional evidence of the effectiveness of our miRNA-based screening protocol, it should be noted that Eip93f, the only other transcription factor whose loss-of-function eliminates Or47b expression, appears on the candidate target lists of two miRNAs whose over-expression eliminates expression of the Or47b reporter. This means we could have identified it in a second round of gene-specific RNAi screening had the list of candidate targets appearing on three lists failed to identify a true positive.

### Validation of a role for Atf3 in Or47b expression

Atf3 is a member of the basic leucine zipper (bZIP) family of transcription factors. In *Drosophila*, Atf3 functions in immune and metabolic homeostasis as well as abdominal morphogenesis (Rynes et al., 2012; Sekyrova et al., 2009). To determine whether Atf3 is expressed in the adult antenna, we amplified and cloned an 800 bp fragment of the *atf3* cDNA from a cDNA library generated by reverse transcription of antennal RNA isolated from wild type Canton S antennae. We used this cDNA as a template for the *in vitro* transcription of an anti-*atf3* riboprobe, and performed RNA *in situ* hybridization on antennal sections of control flies (i.e., the w1118 genetic background strain). We observed that Atf3 is expressed in olfactory neurons all over the antenna, including the distolateral portion where the Or47b-expressing neurons are located (Fig. 4A). We next confirmed that Atf3 is expressed in Or47b neurons by combining an Atf3 protein fusion to eGFP under the transcriptional control of the Atf3 promoter with an Or47b-GAL4 line driving the red fluorescent marker UAS-TdTomato (Fig. 4B).

**Figure 4.**
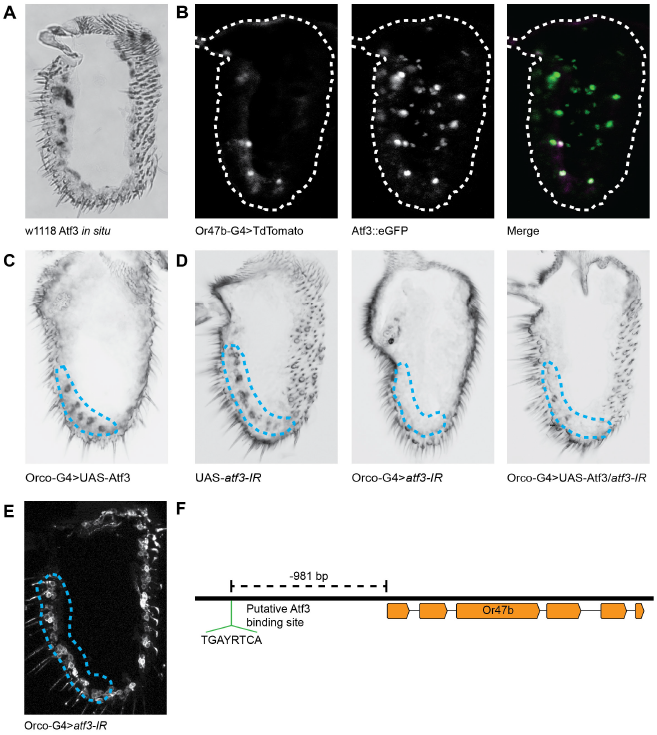
Atf3 is required for the expression of Or47b. A. RNA *in situ* hybridization using an anti-*atf3* probe on control antennae reveals broad expression in adult olfactory neurons. **B**. An Atf3::eGFP protein fusion under the control of the Atf3 promoter is co-expressed with Or47b-GAL4 driving UAS-TdTomato in Or47b neurons. **C**. Over-expression of Atf3 in most adult OSNs using Orco-GAL4 does not expand the expression of Or47b. **D**. When compared to the UAS-*atf3*-IR control (left), Orco-GAL4 driven knockdown of Atf3 eliminates Or47b expression (center), but this can be partially rescued by UAS-Atf3 (right). **E**. Orco-GAL4 driven knockdown of Atf3 does not affect the morphology of the Or47b neurons, nor does it affect Orco expression or dendritic localization. **F**. The Or47b promoter contains a putative Atf3 binding site.

The fact that the other transcription factors implicated in OR expression (e.g., Acj6, Pdm3, Xbp1, Eip93f, etc.) work together as part of a combinatorial code that includes both transcriptional activators and repressors (Jafari et al., 2012), suggests that broad antennal over-expression of Atf3 may expand the expression pattern of Or47b. On the contrary, the combination of UAS-Atf3 with the olfactory coreceptor Orco-GAL4 line, which drives expression in most antennal OSNs, has no effect on the Or47b expression pattern (Fig. 4C). This suggests that either other OSN populations express sufficient repressive transcription factors to prevent Or47b expression or lack an Atf3 binding partner required for Or47b activation.

We used Pbl-GAL4 in our two-tiered miRNA-based screen because it drives expression in peripheral sensory neurons including those of the developing antenna beginning 12-18 hours after puparium formation (APF) (Sweeney et al., 2007). Since Pbl-GAL4 expression begins soon after the birth of the OSNs and long before the earliest OR expression begins at 50 hours APF, it is possible that Atf3 plays a developmental role in Or47b-expressing neurons. Thus, we repeated the Atf3 loss-of-function experiment using Orco-GAL4, which begins to drive expression in antennal OSNs at roughly 80 hours APF, after olfactory development is complete (Larsson et al., 2004). Like the result with Pbl-GAL4, knockdown of Atf3 using Orco-GAL4 eliminates Or47b expression, and this loss of Or47b expression can be partially rescued by re-introduction of Atf3 (Fig. 4D). Consistent with a role for Atf3 in Or47b expression rather than olfactory neurogenesis, knockdown of Atf3 does not affect OSN morphology, Orco expression, or Orco localization (Fig. 4E).

Despite providing ample evidence of a role for Atf3 in Or47b expression, it remains unclear whether Atf3 is acting directly on the Or47b promoter. Miller and Carlson generated a series of GAL4 lines from truncated Or47b promoters. They observed that promoters of 7.6 kilobases down to 419 base pairs drive expression in a small population of neurons in the distolateral antenna similar to the RNA *in situ* pattern circled in blue in Fig. 3F. Further truncation expands the range of labeled cells into the proximal antenna and the maxillary palps (Miller and Carlson, 2010). Their results suggest the existence of an essential repressor binding site in the Or47b promoter between -419 and -342 bp from the transcription start site (TSS) and an essential activator binding site between -219 bp and -119 bp from the TSS. Unfortunately, there is no clear binding site in the Or47b promoter that matches the only published (Sekyrova et al., 2009) consensus sequence recognized by *Drosophila* Atf3, TGACGTCA. There is a published (Rynes et al., 2012) consensus site that would be recognized by mammalian Atf3 (TGAYRTCA) 918 bp upstream of the TSS (Fig. 4F), but this falls outside of the -219 to -119 bp range suggested by Miller and Carlson (Miller and Carlson, 2010). Thus, although further experiments are necessary to determine the exact mechanism of action by which Atf3 regulates Or47b expression, our accelerated miRNA-based screening system permitted the rapid identification of a novel gene involved in Or expression in less than 180 crosses.

## Discussion

Here we report our development of a two-tiered, miRNA-based screening system designed to accelerate the process of genetic screening in *Drosophila*. We generated a collection of transgenic fly stocks that permit the tissue-specific over-expression of miRNAs. Since miRNAs down-regulate their targets, we reasoned that miRNAs could be used for a pooling pre-screen that points the way to a much smaller list of RNAi stocks for follow-up screening than would normally be necessary. We demonstrated this screening method in the olfactory system where we identified a novel role for the transcription factor Atf3 in the expression of the socially relevant odorant receptor Or47b.

Our UAS-miRNA lines are not the first, but previous collections were used for studying miRNA biology directly or in traditional modifier screens (Szuplewski et al., 2012; Bejarano et al., 2012; Schertel et al., 2012). In these cases, it was advantageous to produce different types of insertions (i.e., P-element based, and site-specific) each giving a different level of expression. Since we were interested in using our UAS-miRNA lines for genetic screening, we used PhiC31-mediated site-specific recombination to insert all of our UAS-miRNA lines in the same location on chromosome 2. This simplifies the genetics of a screen and permits comparisons between lines.

The clear advantage to a two-tiered miRNA-based screening strategy is a reduction in the total number of crosses necessary to identify a gene associated with a specific phenotype. Genetic screening always depends on a limited human resource—the patience and endurance of the researcher. In traditional genetic screens the assay for the screened phenotype must be as simple and streamlined as possible to minimize researcher fatigue. In our proof-of-principle screen in the olfactory system, we were able to identify Atf3 in less than 180 crosses. With this level of acceleration, it becomes feasible to design screens with more complicated or time-consuming assays.

It has not escaped our attention, however, that miRNA-based screening is inherently biased and will not be suitable for screening in every tissue or for every phenotype. One type of bias stems from our strict use of endogenous *Drosophila* miRNAs rather than designing short synthetic miRNAs that can target a wider range of genes. Even with a collection of synthetic miRNAs, though, any form of miRNA-based screening will be biased toward identifying genes with longer open reading frames and longer 3’-UTRs. Another potential problem with a miRNA-based approach is the case where a single miRNA produces a synthetic phenotype that is not attributable to a single or predominant target mRNA. Still, we hope that the miRNA-based screening method we describe here will be another useful addition to the geneticist’s toolbox.

## Materials and methods

### UAS-miRNA stock generation

We obtained 44 UAS-DsRed-miRNA constructs from the *Drosophila* RNAi Screening Center originally published by Silver et al. (Silver et al., 2007). We inserted a minimal attB site into these vectors to make them suitable for site-specific insertion into the fly genome. For the rest of the UAS-miRNA stocks, we modified the pKC27-derived attB-containing SST13 UAS vector, which makes use of the split-white system to reduce the time spent on off-target insertions (Yapici et al., 2008). We inserted additional restriction sites between the KpnI and XbaI sites of the SST13 polylinker. Inspired by the design of the Silver et al. UAS-miRNA vectors (Silver et al., 2007), we inserted either the DsRed or mCherry coding sequence downstream of a 5X UAS sequence and renamed these vectors pSS-DsRed and pSS-mCherry (Map and sequence available upon request). Next, we amplified putative pri-miRNA sequences consisting of roughly 400-500 bp surrounding each mature miRNA from Canton S or w1118 genomic DNA and inserted them into the 3’-UTR of the fluorescent marker in each pSS vector. We made constructs for individual miRNAs where possible, but cloned tightly clustered miRNAs en masse. We validated all of our UAS-miRNA vectors with restriction digests and sequencing. Supplemental table 1 summarizes the UAS-miRNA constructs and the primers used to make them. All UAS-miRNA constructs were injected into embryos of the attP-72A landing site strain (n.b., 2nd chromosome) using standard microinjection techniques.

### *Drosophila* stocks

All stocks were maintained at 25°C on conventional media. For validating the UAS-miRNA library, virgin females from GMR-GAL4 (BDSC 9146) and nubbin-GAL4 (BDSC 25754) were mated to UAS-miRNA males. For the two screening steps, virgin females (Pbl-GAL4, UAS-Dcr2;Or47b*>*CD8::GFP, Or92a*>*CD8::GFP) or Orco-GAL4 (BDSC 26818), were mated to either UAS-miRNA or UAS-IR (inverted repeat) males. The crosses were maintained at 25°C until being shifted to 27°C during pupation to enhance GAL4 expression. All UAS-miRNA and UAS-IR lines that reduce the expression of the GFP reporters were checked 2-3 times to ensure reproducibility. Two different UAS-Atf3-IR lines (BDSC 26741 and VDRC 105036) produced identical phenotypes.

### Immunostaining

Brains were dissected and stained as previously described (Laissue et al., 1999) 3-4 days after eclosion with an anti-GFP antibody (1:1000, Molecular Probes) and the monoclonal antibody nc82 (1:50, DHSB), which recognizes neuropil. Frozen antennal sections were stained as described previously (Larsson et al., 2004) using an anti-GFP antibody (1:1000, Molecular Probes) and an anti-Orco antibody (1:5000) raised against the peptide (SSIPVEIPRLPIKS) by AbFrontier.

### *In situ* hybridization

*In situ* hybridization on frozen antennal sections was carried out as previously described (Vosshall et al., 1999) using probes generated by *in vitro* transcription from cDNAs for Or47b, Or92a, and Atf3 in the pGEMT-Easy vector (Promega). The cDNAs were cloned using standard methods using the following primers: Or47b (CGGGTTATCAATCAAATCTCAGCC, GTGGAACCTCTTATCACTGACCTC), Or92a (TGTGGTGGGCGAAATCGCGT, TGTGGTGCGGAGCAGTGCAA), and Atf3 (GTTCAATTCCAACATACCGGCC, GATTTCAGCATGTCCACCAACTTT). Or47b/Or92a *in situs* were developed for 3 hours, but Atf3 *in situs* were developed for 3 days.

## Author contributions

W.D.J. developed the idea for miRNA-based screening. S.B., M.S., SY.B., and Y.K. generated and tested the UAS-miRNA fly stocks. S.B. carried out the screen and performed the rest of the experiments. S.B. and W.D.J. analyzed the data and wrote the paper.

## Acknowledgements

We thank Mattias Alenius (Linköping, Sweden) for providing the Pbl-GAL4,UAS-Dcr2;Or47b*>*CD8::GFP, Or92a*>*CD8:GFP stock, Mirka Uhlirova (Köln, Germany) for providing the UAS-Atf3 and Atf3::eGFP stocks, as well as the VDRC and TRiP at Harvard Medical School (NIH/NIGMS R01-GM084947) for providing transgenic RNAi fly stocks. This work was supported by grants to W.D.J. (2010-0006217 and 2013R1A1A2011339) from the National Research Foundation of Korea (NRF), which is funded by the Ministry of Science, ICT and Future Planning (MSIP), Republic of Korea.

